# Boosting N-terminally anchored yeast surface display via structural insights into *S. cerevisiae* Pir proteins

**DOI:** 10.1101/2023.04.25.538238

**Authors:** Tea Martinić Cezar, Mateja Lozančić, Ana Novačić, Ana Matičević, Dominik Matijević, Beatrice Vallee, Vladimir Mrša, Renata Teparić, Bojan Žunar

**Affiliations:** Laboratory for Biochemistry; Department of Chemistry and Biochemistry, Faculty of Food Technology and Biotechnology, University of Zagreb, Pierottijeva 6, 10000 Zagreb, Croatia; Centre de Biophysique Moléculaire (CBM), CNRS, UPR 4301, University of Orléans and INSERM, 45071 Orléans Cedex 2, France

**Keywords:** surface display, Pir proteins, Hsp150, cell wall, *Saccharomyces cerevisiae*

## Abstract

Surface display co-opts yeast’s innate ability to embellish its cell wall with mannoproteins, thus converting the yeast’s outer surface into a growing and self-sustaining catalyst. However, the efficient toolbox for converting the enzyme of interest into its surface-displayed isoform is currently lacking, especially if the isoform needs to be anchored to the cell wall near the isoform’s N-terminus. Aiming to advance such N-terminally anchored surface display, we employed *in silico* and machine-learning strategies to study the 3D structure, function, genomic organisation, and evolution of the Pir protein family, whose members evolved to covalently attach themselves near their N-terminus to the β-1,3-glucan of the cell wall. Through the newly-gained insights, we rationally engineered 14 *S. cerevisiae* Hsp150 (Pir2)-based fusion proteins. We quantified their performance, uncovering guidelines for efficient yeast surface display while developing a construct that promoted a 2.5-fold more efficient display than the full-length Hsp150 and a Pir-tag, i.e., a peptide spanning only 4.5 kDa but promoting as efficient surface display as the full-length Hsp150. These constructs fortify the existing surface display toolbox, allowing for a prompt and routine refitting of any protein into its N-terminally anchored isoform.

**Graphical abstract:** 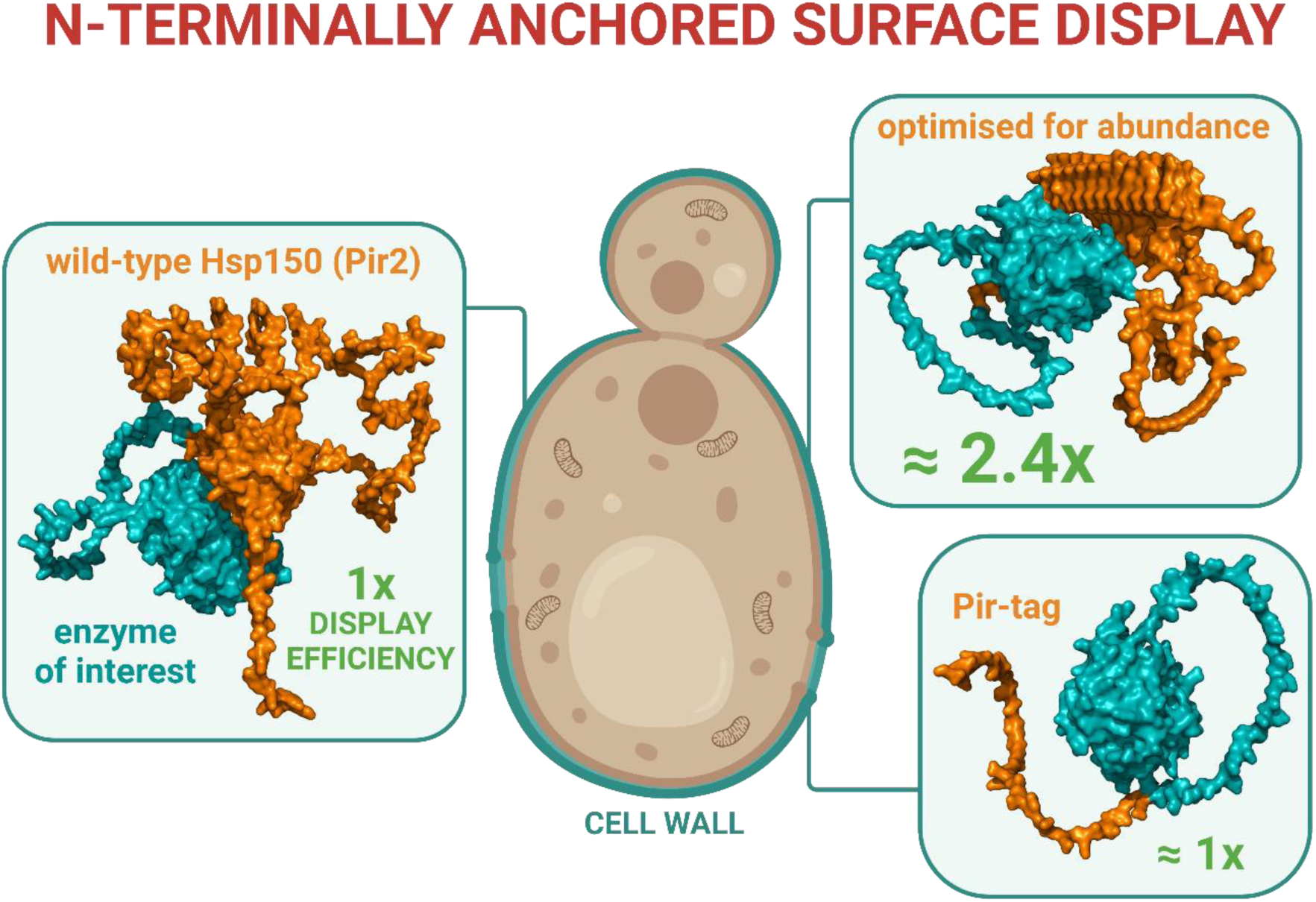

## 1. Introduction

Unlike Metazoan cells, microbes encase themselves in a cell wall, a protective layer through which they interact with the environment. While they build this structure primarily from polysaccharides, they also modify it heavily, purposefully meshing it with other macromolecules to adapt its properties and function (Klis et al., 2002; Orlean, 1988). Yeast cells, in particular, weave their vegetative cell wall from an inner, polysaccharide-based layer, to which they covalently and noncovalently attach mannoproteins, thus moulding the cell wall’s outer layer (Cabib et al., 1982; Orlean, 2012).

Being naturally malleable, the yeast cell wall can, in principle, be modified for many biotechnological applications. By re-engineering mannoprotein synthesis, i.e., displaying enzymes of interest on the cell’s surface, we could convert the cell wall’s outer layer into a growing and self-renewing catalytic surface, a living material (Molinari et al., 2021; Nguyen et al., 2018; Žunar et al., 2022a). Such a surface is ideal for catalysing reactions reliant on toxic, highly-concentrated, or cell-impermeable substrates, while obtaining readily purifiable products of enzymatic reactions (Han et al., 2018; Teymennet-Ramírez et al., 2022).

The most promising approach to cell surface engineering relies on surface display (Gai and Wittrup, 2007; Lozančić et al., 2019; Tanaka et al., 2012; Ueda, 2016). This protein engineering technique reconfigures intracellular proteins for secretion and cell wall binding, commonly by fusing them with native cell wall proteins while respecting the mechanisms that bind native proteins to the cell wall’s inner, polysaccharide layer (Abe et al., 2004; Boder and Wittrup, 1997; Matsumoto et al., 2002; Nakamura et al., 2001; Van der Vaart et al., 1997). To bind to the cell wall covalently, the enzyme of interest is fused to either a GPI-anchored cell wall protein or a Pir protein, at its C- or N-terminus, respectively. However, some enzymes tolerate fusions with native cell wall proteins at only one of their termini, otherwise failing to fold properly, to pass quality control in the secretory pathway, or to be flexible enough to catalyse the intended reaction (Teparic and Mrsa, 2015; Teymennet-Ramírez et al., 2022; Wang et al., 2005). Thus, only a subset of theoretically possible fusion proteins performs well *in vivo*, which hinders the full-scale application of surface display.

Pir proteins (<u>p</u>roteins with internal repeats) are a family of five *Saccharomyces cerevisiae* proteins (Pir1, Hsp150 (Pir2), Pir3, Cis3 (Pir4), and Pir5) whose sequence features are well documented. All five Pir proteins are directed to the secretory pathway through their N-terminal signal peptides, which are followed by motifs of 18 amino acid residues, after which they were named and which they encode one to eleven times, repeating them *in tandem*. Through these repeats, Pir proteins bind to the cell surface covalently, with the γ-carboxyl group of the second glutamate within the repeat binding to β-1,3-glucan of the inner polysaccharide layer (Ecker et al., 2006). Following the repeat region, Pir proteins encode a well-conserved C-terminal region that potentially binds the cell wall through disulphide bonds (Castillo et al., 2003).

Less is known about the structure and function of Pir proteins. Pir1-Pir4 are highly O-glycosylated and important for cell wall stability, as their quadruple deletion mutant grows slowly, has aberrant morphology, and is sensitive to cell wall-stressing compounds (Mrša and Tanner, 1999). Furthermore, Pir proteins are distinctly expressed throughout the cell cycle and under stress conditions (Jung and Levin, 1999; Spellman et al., 1998), with Pir5 being expressed only during meiotic development (Enyenihi and Saunders, 2003).

Pir proteins can be used as anchors for surface display (Yang et al., 2014). However, previously employed strategies relied only on known sequence features and native restriction sites. Moreover, they could display only smaller proteins, as the proteins of interest had to be anchored to the near-full-length Pir protein. Hence, further innovation of N-terminally anchored surface display was hampered by the unfamiliar 3D structure and ambiguous function of Pir proteins.

Aiming to enhance N-terminally anchored yeast surface display, we focused on essential properties of the Pir protein family, i.e., their 3D structure and putative function, genomic organisation, and the number and conservation of Pir repeats. We explored these topics through modern genomic, structural, and evolutionary *in silico* approaches and machine-learning strategies. Using these newly-gained insights, we rationally engineered *S. cerevisiae* Hsp150 (Pir2), developing constructs and guidelines for highly efficient and easily implementable N-terminally anchored yeast surface display.

## 2. Methods

### 2.1. Predicting and exploring protein structures and conformation switching

Protein sequences were aligned with NBCI’s blastp (Johnson et al., 2008), using standard parameters. For Pir1-Pir5, tertiary structures and the corresponding structural confidence scores were obtained from EMBL-EBI AlphaFold Protein Structure Database (Jumper et al., 2021; Varadi et al., 2022). The structures were visualised in PyMOL 2.5 (Schrodinger, 2022) and structurally superposed with PyMOL’s align command. The structural similarity of the β-trefoil domain was investigated with Dali (Holm, 2020), using a heuristic PDB search. The structures of the identified structurally similar β-trefoil domains were superposed with PyMOL’s cealign command. The structures of rationally engineered Hsp150-β-lactamase fusion proteins were predicted with Colabfold (Mirdita et al., 2022), a version of the deep neural network Alphafold2 coupled with Google Collaboratory, using default parameters.

To model the Hsp150 (Pir2) conformation-switching, we emulated Wayment-Steele et al. (2022), i.e., we clustered the Colabfold-generated multiple sequence alignment and specified each alignment branch as a separate Colabfold input.

### 2.2. Investigating Pir genomic loci and phylogeny

Homologous genomic loci within the *Saccharomyces* clade were identified by the Smith-Waterman alignment of *S. cerevisiae* Pir regions and whole genome sequences of the neighbouring *Saccharomyces* species. The analysis was performed in Geneious Prime (Dotmatics, 2022) and the results manually curated, with the number of Pir repeats counted by HMMER (Finn et al., 2011).

Pir phylogeny was inferred in MEGA 11 (Tamura et al., 2021) by applying to the 406 Pir homologues identified in Lozančić et al. (2021) the maximum likelihood method and Dayhoff matrix-based model (Schwarz and Dayhoff, 1979). Positions with less than 95% site coverage were eliminated. An initial heuristic search was performed with neighbour-join and BioNJ algorithms, and evolutionary rate differences among sites were modelled with a discrete Gamma distribution (5 categories, parameter = 2.3669).

The Ykl162c-a signal sequence was analysed with specialised web-hosted machine learning methods DeepSig (Savojardo et al., 2018) and SignalP (Teufel et al., 2022). Gene expression heatmaps were obtained from Saccharomyces Genomics Viewer (http://sgv.genouest.org/) (Lardenois et al., 2011).

### 2.3. Media and growth conditions

*E. coli* were grown overnight at 37 °C at either 200 rpm in liquid 2xYT media (16.0 g/l tryptone, 10.0 g/l yeast extract, 5.0 g/l NaCl) or on LB plates (10.0 g/l tryptone, 5.0 g/l yeast extract, 5.0 g/l NaCl, 15.0 g/l agar), supplemented with 100 µg/ml of ampicillin. Wild-type *S. cerevisiae* BY 4741 was grown at 30 °C/180 rpm in standard YPD medium (20.0 g/l peptone, 10.0 g/l yeast extract, 20.0 g/l glucose, and 20.0 g/l agar for solid media). *S. cerevisiae* strains carrying reporter plasmids were grown at 30 °C/180 rpm in chemically defined media without histidine (-his: 6.70 g/l Difco Yeast nitrogen base without amino acids, 20.0 g/l glucose, 1.6 g/l drop-out without histidine, and 20.0 g/l agar for solid media). To induce the *PHO5* promoter, yeast cells were first grown in a repressive and then in a permissive medium, i.e., in -his medium supplemented with 1 g/l of KH_2_PO_4_ and then in -his medium lacking all phosphate sources, respectively, as in Novačić et al. (2022).

### 2.4. Plasmid and strain construction

Plasmids were constructed in NEB Stable *E. coli* using restriction cloning, Q5 polymerase, and HiFi Assembly (New England Biolabs, Frankfurt am Main, Germany), according to the manufacturer’s instructions. *S. cerevisiae* BY 4741 was transformed as in Gietz and Schiestl (2007). The details of the constructions are provided in the Supplementary material. Plasmid DNA was isolated with NucleoSpin Plasmid Mini Kit and extracted from an agarose gel with NucleoSpin Gel and PCR Clean up Kit (Macherey-Nagel, Duren, Germany). All constructs were verified by restriction digest and Sanger sequencing. The results of the plasmid analyses agreed with computed restriction maps. Primer synthesis was outsourced to Macrogen Europe (Amsterdam, Netherlands), *de novo* gene synthesis to Genewiz (Leipzig, Germany), and Sanger sequencing to Microsynth (Balgach, Switzerland).

### 2.5. Isolation of cell wall proteins

Cell wall proteins were isolated using the same methodology as in Lozančić et al. (2021). In short, cells were grown in the -his medium without phosphate until the stationary phase, resuspended in the potassium phosphate (KP) buffer (50 mM, pH 8), mechanically disrupted with glass beads, and washed with KP buffer four times. Non-covalently bound proteins were isolated by boiling the cell wall fraction twice in the reducing Laemmli buffer (0.0625 M Tris, pH 6.8, 2% SDS, 5% v/v β-mercaptoethanol, 0.001% bromophenol blue), which extracted non-covalently bound proteins to the supernatant. To isolate Pir proteins bound to the cell wall covalently, the cell walls stripped of the non-covalently bound proteins, contained within the solid fraction from the previous step, were washed four times with KP buffer, twice with deionised water, and incubated overnight at 4 °C in 30 mM NaOH. Protein samples were standardised by normalising the amount of the extraction buffer and extract used for electrophoresis to the mass of wet cell walls, which was measured before stripping non-covalently bound proteins from the cell walls.

### 2.6. Immunoblotting

Isolated proteins were resolved on 10% SDS-PAGE gels, transferred to a nitrocellulose membrane by semidry transfer (Trans-Blot TurboTM Transfer System, BioRad, Hercules, United States), and analysed by immunoblotting using standard procedures, as in Novačić et al. (2021). Blots were developed with Clarity Western ECL substrates (Biorad, Hercules, United States) and visualised with a C-DiGit Blot scanner (LI-COR Biosciences, Lincoln, United States).

For Ykl162c-a experiments, strains were grown to the exponential (OD_600_ 2) phase in the YPD medium, and total proteins extracted as in Kushnirov (2000). Myc-tagged Ykl162c-a was probed with anti-c-Myc (1:1,000 dilution; 9E10, Santa Cruz Biotechnology, Dallas, United States) as a primary and mouse IgG κ-binding protein horseradish peroxidase (1:50,000 dilution; Santa Cruz Biotechnology, Dallas, United States) as a secondary antibody. For Hsp150-β-lactamase experiments, isolated cell wall proteins were probed with anti-HA peroxidase-conjugated antibody (1:1,250 dilution; 11667475001, Roche, Basel, Switzerland).

### 2.7. Measuring β-lactamase activity

Cells were inoculated into 5 ml of -his medium supplemented with KH_2_PO_4_ and grown overnight at 30 °C/180 rpm. The following day, stationary phase cells were diluted to 0.5 OD_600_/ml in 15 ml -his medium supplemented with KH_2_PO_4_ and incubated at 30 °C/180 rpm until reaching 2 OD_600_/ml. To induce the *PHO5* promoter, cells were washed with sterile deionised water (sdH_2_O), diluted to 0.3 OD_600_/ml in 15 ml of -his medium without phosphate, and grown overnight at 30 °C/180 rpm, thus reaching 2 OD_600_/ml in the morning. Such cells were washed in sdH_2_O, then in potassium phosphate (KP) buffer (pH 7, 50 mM), and resuspended in the same buffer to 100 OD_600_/ml. Next, 1 OD_600_ of cells was resuspended in 475 μl of KP buffer, thermostated at 30 °C/1,200 rpm/2 min, mixed with 25 μl of nitrocefin (1 mM, dissolved in 50 mM KP buffer with 5% DMSO, pH 7), and incubated at 30 °C/1,200 rpm/5 min. The reaction was stopped by centrifuging the cells (8,000 rpm/30 s) and measuring A_482_ of the supernatant. The activity of each strain was measured in technical triplicates and at least biological duplicates.

### 2.8. Yeast staining, confocal microscopy, and flow cytometry

The yeast culture was grown to the exponential phase (OD_600_ 2), centrifuged (13,000 rpm/30 s), washed with PBS (137 mM NaCl, 2.7 mM KCl, 10 mM Na_2_HPO_4_, 1.8 mM KH_2_PO_4_, pH 7.4), and cells resuspended in either 100 μg/ml of concanavalin A-FITC (to visualise mannoproteins; C7642, Sigma-Aldrich, St. Louis, United States), 1,000 μg/ml of calcofluor white (to visualise chitin; 18909, Sigma-Aldrich, St. Louis, United States), or 500 μl/ml of aniline blue (to visualise β-glucans; 415049, Sigma-Aldrich, St. Louis, United States) as in Okada and Ohya (2016). The staining proceeded for 10 min. Each experiment was performed in triplicate.

After staining, the suspension was mixed with 3 ml of PBS and analysed by flow cytometry. For each condition, BD LSRFortessa X-20 (Franklin Lakes, Unite States) recorded 50,000 events, using a 488 nm laser and 530/30 nm emission filter for concanavalin A-FITC, a 405 nm laser and 450/50 nm emission filter for calcofluor white, or a 405 nm laser and 525/50 nm emission filter for aniline blue. Data were analysed in the R computing environment (R Core Team, 2022) using openCyto (Finak et al., 2014) and ggcyto packages (Van et al., 2018), with the fluorescence assessed on the flowJo biexponential scale.

For confocal microscopy, following staining, the cells were centrifuged (13,000 g/30 sec), resuspended in 10 μl of PBS and imaged on a ZEISS LSM 980 confocal microscope (Carl Zeiss, Jena, Germany) under a 40x oil objective, using laser wavelengths of 488 and 561 nm, as in Žunar et al. (2022b). Images were processed using Zeiss Zen Blue software.

To measure supernatant fluorescence, cells were grown in 10 ml of standard -ura medium to stationary phase, at 30 °C/180 rpm/overnight, and centrifuged (3,000 rpm/5 min). The red fluorescence was measured with a Perkin Elmer LS55 fluorescence spectrometer (Perkin Elmer, Waltham, United States), by exciting 3 ml of supernatant in quartz cuvettes at 569 nm and measuring emission at 594 nm.

## 3. Results

### 3.1. *S. cerevisiae* Pir proteins encode lectin-like β-trefoil domains

Aiming to advance the N-terminally anchored Pir-based surface display, we first sought more insight into the 3D structures of native *S. cerevisiae* Pir proteins. *S. cerevisiae* encodes five Pir proteins, Pir1, Hsp150 (Pir2), Pir3, Cis3 (Pir4), and Pir5, each with a unique number of Pir repeats (Figure 1A). All five Pir proteins are encoded in just two homologous loci, which arose by whole genome duplication, approximately 100 million years ago (Kellis et al., 2004). As such, Pir1 and Pir5 form one, and Pir3 and Hsp150 another paralogue pair. Interestingly, while Cis3 does not have a genuine paralogue pair, homologous locus encodes Ykl162c-a, a 50-amino-acid peptide annotated as a dubious open reading frame unlikely to encode a functional protein. However, the final 35 residues of Ykl162c-a are homologous to the Cis3 C-terminus.

**Figure 1:**
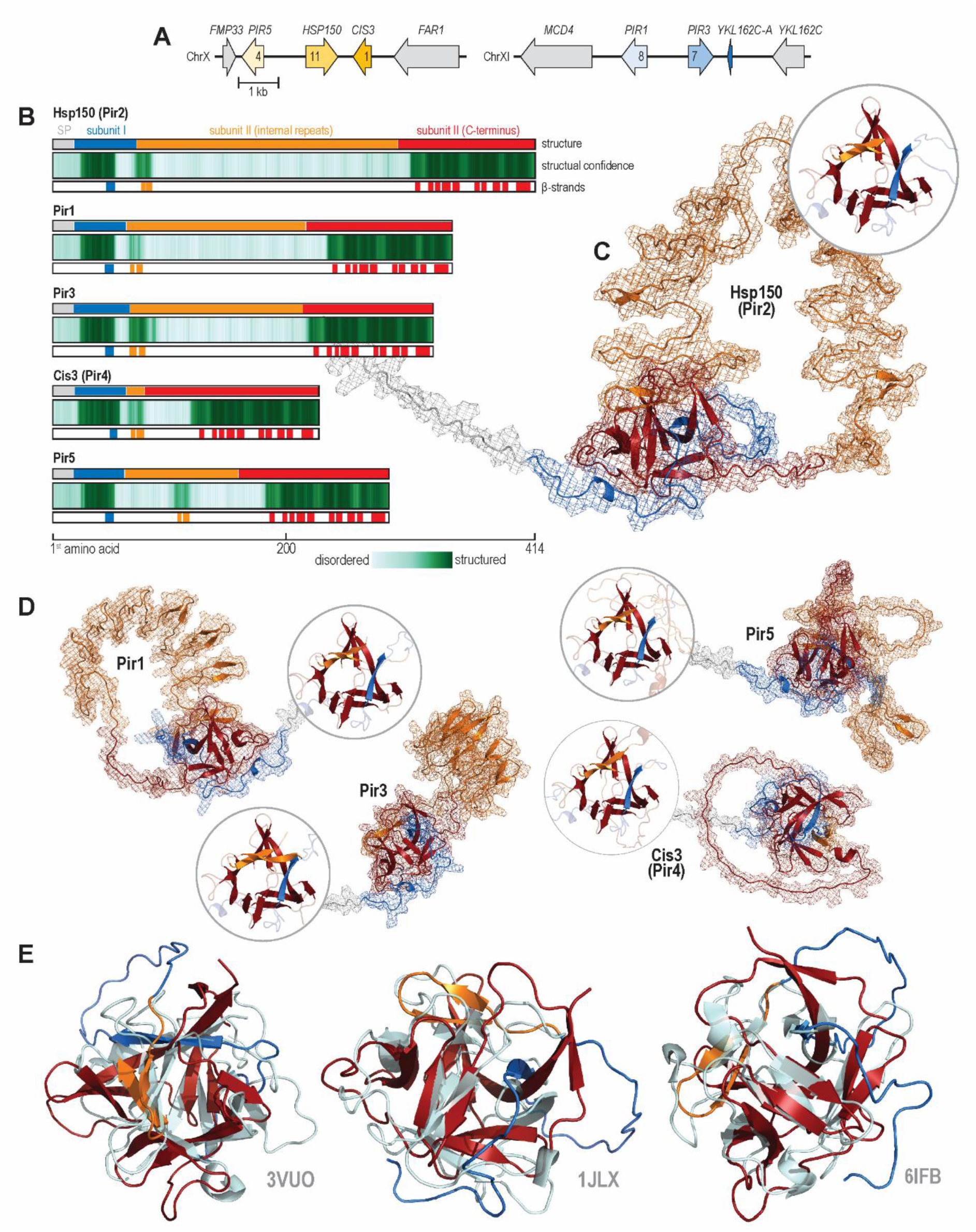
*S. cerevisiae* Pir proteins encode lectin-like β-trefoil domains. A) *S. cerevisiae* Pir proteins are encoded by the two homologous loci on chromosomes X and XI. Arrows denote gene orientation and the numbers within them the number of Pir repeats. B) The protein sequences of five *S. cerevisiae* Pir proteins (upper track) are homologous, encoding a signal peptide (SP, grey), a small subunit I (blue), and a large subunit II. Subunit II consists of an N-terminal part carrying internal Pir repeats (yellow) and a well-conserved C-terminal part (red). For each Pir protein, Alphafold2 models only part of the subunit I, one internal repeat, and the C-terminus of subunit II as well structured (middle track, dark green) while suggesting that the remaining parts of the proteins are intrinsically unstructured (faint green). Moreover, Alphafold2 predicts that well-structured parts of the proteins encode thirteen β-strands (lower track). C) Alphafold2-structural model of Hsp150 (Pir2), an archetypal Pir protein. Hsp150 is a knotted protein with a single well-structured β-trefoil domain made of thirteen β-strands. The colour scheme follows panel A. The details of the β-trefoil domain are presented in the circular inset, which plots only β-strands in full colour. D) Predicted structures of the remaining four *S. cerevisiae* Pir proteins. Circular insets show structurally-aligned β-trefoil domains. E) PyMOL’s structural alignment of Hsp150 β-trefoil domain and several structurally similar crystallised proteins, as determined by Dali. The panels show structural alignments between the Hsp150 domain and nontoxic non-hemagglutinin subcomponent (NTNHA) from *Clostridium botulinum* (PDB: 3VUO), agglutinin from *Amaranthus caudatus* (PDB: 1JLX), and β-trefoil lectin from *Entamoeba histolytica* (PDB: 6IFB). The blue-orange-red colour scheme of the Hsp150 domain follows panel A. The aligned PDB structures are shown in grey.

At the amino acid sequence level, all five *S. cerevisiae* Pir proteins are homologous to each other (Figure 1B). Each protein encodes a signal peptide, a subunit I, a Kex2-processing site (Lys-Arg) at which Kex2 cleaves the Pir protein into two subunits that remain non-covalently bound to each other, and a subunit II, with the subunit terminology being initially developed to describe the Hsp150 features (Russo et al., 1992). Subunit II can be further divided into an N-terminal region, i.e., the internal repeat-encoding region of variable length, and a well-conserved C-terminal region.

To infer the 3D structural features of Pir proteins, we examined their Alphafold2-predicted models. For each Pir homologue, Alphafold2 suggested that only some regions of the protein form well-structured domains, specifically subunit I, one of the internal repeats, and the C-terminal region of subunit II, which together folded into thirteen β-strands per protein (Figure 1B). Interestingly, one β-strand was always encoded by the subunit I, two by one of the Pir repeats, and ten by the C-terminal end of the subunit II. Conversely, Alphafold2 suggested that the remaining parts of the proteins, i.e., signal peptide and all-but-one internal repeats, were intrinsically disordered.

We closely analysed the Alphafold2-predicted structure of Hsp150, an archetypal Pir protein. Alphafold2 modelled this protein as topologically knotted, with thirteen β-strains folding into a single well-structured β-trefoil domain (Figure 1C). As in other *S. cerevisiae* Pir proteins, β-strains originated from non-adjacent parts of the protein. Moreover, Alphafold2 predicted that the remaining ten Pir repeats form a disordered loop, which started and ended at the β-trefoil domain. Other *S. cerevisiae* Pir proteins followed the same basic structure, forming β-trefoil domains with highly conserved β-stranded cores yet divergent β-trefoil turns (Figure 1D).

To deduce the function of the Pir proteins, we compared the Hsp150-predicted β-strain domain with experimentally solved protein structures available in Uniprot, using Dali (Holm, 2020). This methodology uncovered significant similarities to several lectin β-trefoil domains (Fig 1E), e.g., domains contained in nontoxic non-hemagglutinin subcomponent (NTNHA) from *Clostridium botulinum* (PDB: 3VUO), agglutinin from *Amaranthus caudatus* (PDB: 1JLX), and β-trefoil lectin from *Entamoeba histolytica* (PDB: 6IFB). Thus, Pir proteins are likely a family of topologically knotted cell wall-bound lectin-like proteins.

### 3.2. Pir repeats have the potential to fold into a two-faced β-helical domain

Next, we focused on the most notable feature of Pir proteins, their internal (Pir) repeats. We first traced the evolutionary origin of Pir repeats. The Alphafold2 structural model of Cis3, which carries only one Pir repeat, showed that the repeat folds into two β-sheets that formed an integral part of the β-trefoil domain, suggesting it represents the original Pir repeat. Accordingly, the additional Pir repeats likely arose from the expansion of the original two β-sheets of the β-trefoil domain.

Next, we investigated the evolutionary stability of the repeats. At the DNA level, Pir repeats are encoded as near-perfect tandem repeats and thus are predisposed to be intrinsically evolutionary unstable. An evolutionary stably-maintained repeat number would suggest positive selection and imply additional repeat functionality, given that a single Pir repeat suffices to form the β-trefoil domain and covalently bind the protein to the cell wall (Ecker et al., 2006).

To assess the evolutionary stability of Pir repeats, we first mapped their numbers across the *Saccharomyces* clade. We identified homologous Pir loci in non-laboratory probiotic *S. cerevisiae* var. *boulardii* and the remaining seven *Saccharomyces* species, which diverged up to 17 million years ago (Shen et al., 2020), and we counted the repeats. The analysis showed that each of the five *S. cerevisiae* Pir proteins had a characteristic repeat number, which they maintained with only minor variations (Figure 2A). Interestingly, this analysis also uncovered that Pir6, a lost *S. cerevisiae* Cis3 paralogue, persists in other *Saccharomyces* species.

**Figure 2:**
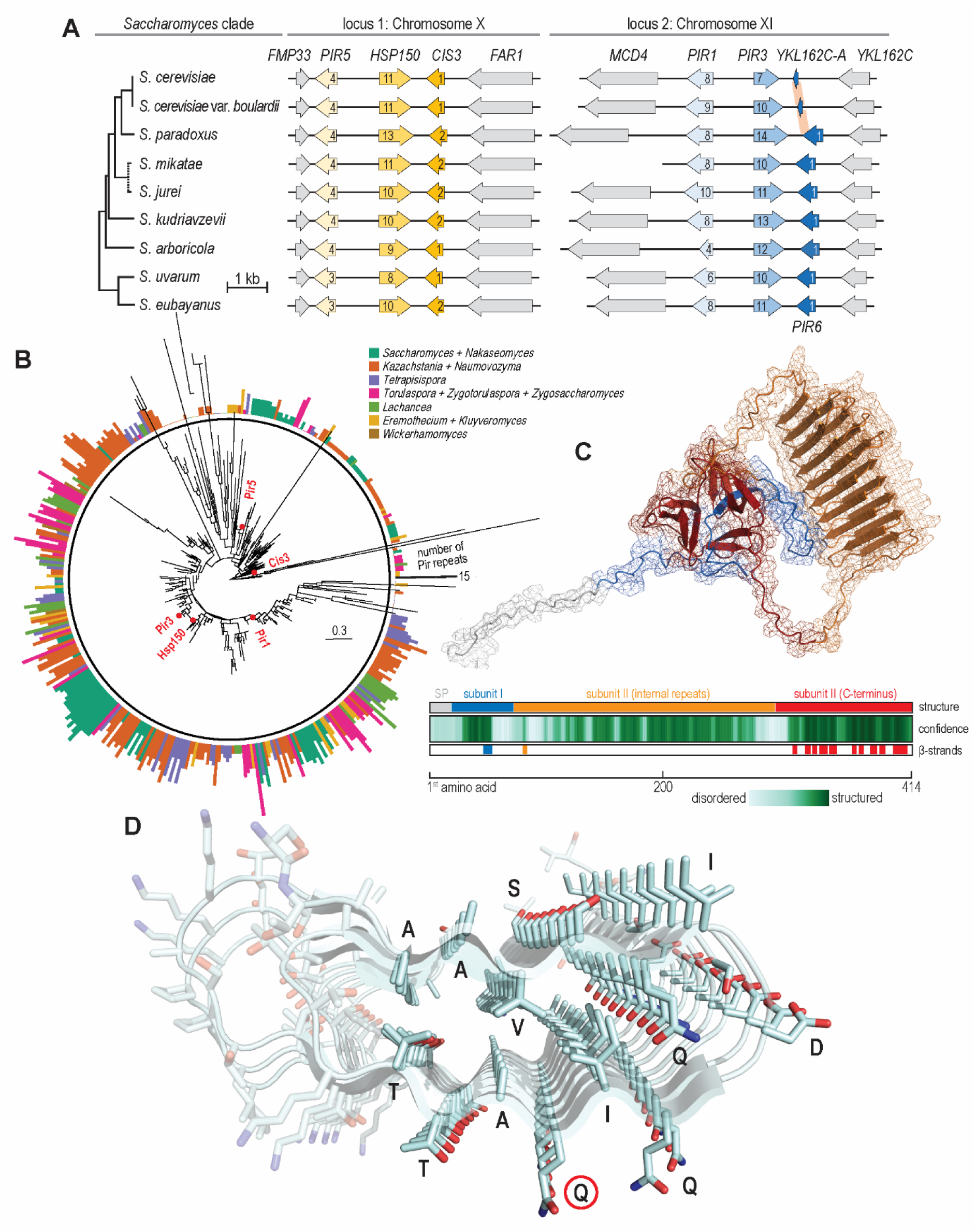
Distribution and putative folding of Pir repeats. A) Homologous Pir loci across the *Saccharomyces* clade. Only *S. cerevisiae* does not encode Pir6, a Cis3 homologue. Arrows denote gene orientation and numbers within them the number of Pir repeats. B) Unrooted phylogenetic tree of 406 Pir homologues across 76 species of budding yeasts. Red points denote the positions of *S. cerevisiae* homologues. The bar height and colour in the outer rim show the number of Pir repeats in each homologue and its clade of origin, respectively. C) Alphafold2 model of Hsp150 (Pir2) with folded Pir repeats. The colour scheme of the structural model and the summary tracks follow Figure 1. D) Sideview of the Hsp150 Pir repeats folded into two-faced β-helix, accentuating ordered folding of amino acid residues. Glutamine residues through which Pir proteins covalently bind to the cell wall are encircled in red.

To test whether Pir repeats remain stably expanded even beyond the *Saccharomyces* clade, we looked at their distribution within predicted proteomes of neighbouring budding yeasts. We analysed 406 Pir proteins from 76 species that diverged from *S. cerevisiae* up to 125 million years ago (Lozančić et al., 2021) (Figure 2B). While the repeat number varied appreciably across the dataset, only Cis3 homologues retained a single Pir repeat. Other Pir homologues consistently maintained more than one Pir repeat, once again suggesting a positive selection for multiple repeats.

Although Alphafold2 modelled all but one internal repeat as an intrinsically disordered loop (Figure 1C), we wondered whether such a disordered structure could nevertheless switch conformation, i.e., fold. Emulating a recent Alphafold2-based study of protein conformation switching (Wayment-Steele et al., 2022), we focused on the *S. cerevisiae* Hsp150, which carries 11 internal repeats. We clustered the Colabfold-generated multiple sequence alignment of Hsp150 and performed the Alphafold2 modelling using each alignment branch as a separate input. Strikingly, the most populated branch (212/288 sequences) produced a 3D structure whose internal repeats stacked one upon other, folding into a well-ordered right-handed, two-faced β-helix (Figure 2C). In such β-helix, each internal repeat formed a well-conserved β-sheet-turn-β-sheet motif. Notably, in such a structure, glutamine residues, through which Pir proteins covalently link to β-1,3-glucan (Ecker et al., 2006), pointed away from the β-helix, theoretically allowing for cell wall binding (Figure 2D). Thus, internal repeats have the potential to fold into a two-faced β-helical domain, through which Pir proteins remain covalently bound to the cell wall.

### 3.3. *S. cerevisiae* encodes a Pir6 remnant

Next, we focused on Ykl162c-a, a seemingly devolved remnant of *S. cerevisiae* Pir6. Ykl162c-a is not specific to the S288c background as it is also present in other commonly used laboratory *S. cerevisiae* strains (Saccharomyces Genome Database, data not shown) and probiotic *S. cerevisiae* var. *boulardii* (Figure 2A). As full-length Pir6 persists in the remaining seven *Saccharomyces* species, including closely related *S. paradoxus*, *S. cerevisiae* must have lost it recently, within the last 5 million years (Shen et al., 2020), probably through frameshift, as the Pir6 loci of *S. cerevisiae* and *S. paradoxus* differ in several indels.

Out of the 50 amino acid residues of Ykl162c, only the final 35 are homologous to the C-terminus of its paralogue Cis3. Unexpectedly, a bioinformatic analysis suggested Ykl162c-a evolved a new signal sequence at its N-terminus (Figure 3A, DeepSig reliability: 1.0, SignalP 6.0 probability: 0.572234, cleavage site between residues 21 and 22). This signal sequence, made of a characteristic hydrophobic core (H-region) flanked by N- and C-regions (Liaci and Förster, 2021), evolved through two single base-pair frameshifts 40 bp apart, with the resulting region encoding 15 novel amino acid residues. To functionally validate the predicted signalling peptide, we Myc-tagged the native, chromosomally-encoded Ykl162c-a at its C-terminus and preceded it with the constitutive *TEF1* promoter. By immunoblotting, we detected such tagged and overexpressed Ykl162c-a in the whole cell extracts (Figure 3B), thus confirming it can be stably expressed. To validate the putative signal sequence, we replaced the C-terminal Myc-tag of the constitutively overexpressed Ykl162c-a with the red fluorescent protein ymScarletI and, by measuring the fluorescence of the medium, followed its secretion. Compared to the control strain, in which ymScarletI was not preceded by Ykl162c-a, the strain with ymScarletI-tagged Ykl162c-a produced a significantly more fluorescent medium (17.7328 ± 1.208898 AU vs 25.0024 ± 5.023018 AU, p <0.03). Thus, Ykl162c-a encodes a novel functional signal sequence.

**Figure 3:**
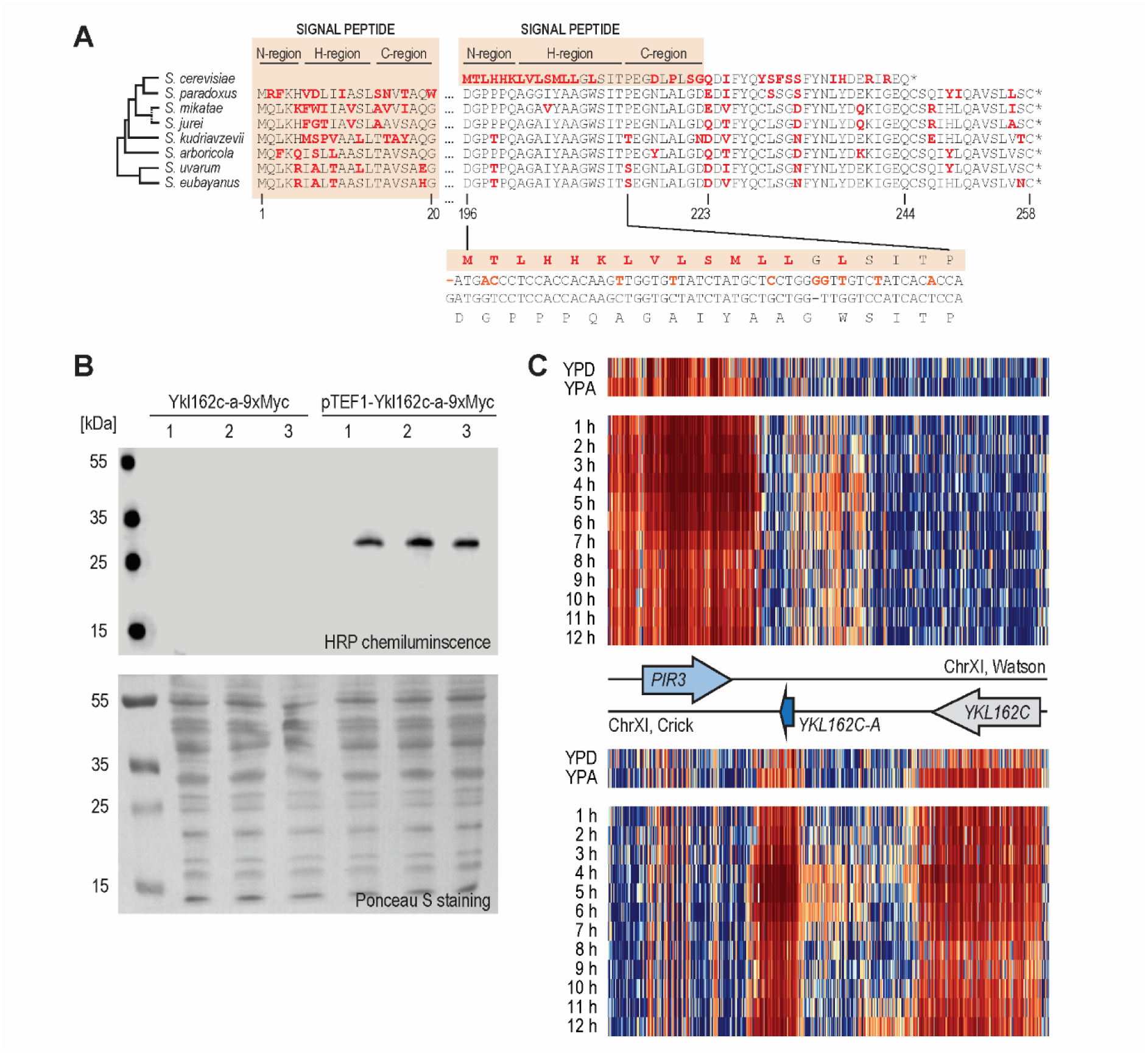
*S. cerevisiae* lacks full-length Pir6, instead encoding 50 aa-long Ykl162c-a. A) An alignment of Ykl162c-a and Pir6 homologues, with red letters indicating mismatches and light orange boxes highlighting signal sequences. The lower inset shows the alignment between *S. cerevisiae* and *S. paradoxus* loci encoding Ykl162c-a, i.e., Pir6 C-terminus, with orange letters indicating mismatches. B) Immunoblot of the C-terminally Myc-tagged Ykl162c-a, expressed from its native promoter (left) and from constitutive strong *TEF1* promoter (right), analysed in biological triplicates from exponentially growing cultures. C) Heatmaps visualising gene expression from the Watson (upper heatmaps) and Crick strands (lower heatmaps) of diploid *S. cerevisiae YKL162c-A* locus show that the locus is expressed in the respiring (YPA) and sporulating (1 h-12 h in SP medium) but not in the fermenting diploid cells (YPD). The colour scale shifts from blue (not expressed) through white to red (strongly expressed).

Moreover, we investigated the expression pattern of native Ykl162c-a. With immunoblotting, we were unable to detect expression of the Myc-tagged Ykl162c-a from its native promoter in the BY 4741 total protein extracts (Figure 3B), despite analysing exponentially-growing fermenting cells (grown in YPD medium to OD_600_ of 0.2, 2, and 4), respiring cells (grown in standard SP and YPA medium), stationary phase cells (grown in YPD medium to OD_600_ of 10), cell wall-stressed cells (grown for 5 h in YPD supplemented with calcofluor white), or heat-shocked cells (grown at 37 °C or 42 °C for 5 h). Furthermore, we were unable to detect secreted Ykl162c-a, despite immunoblotting acetone-precipitated proteins secreted to the medium (data not shown).

However, the Saccharomyces Genomics Viewer (Lardenois et al., 2011) indicated that the *YKL162C-A* locus is transcribed to a potentially significant level in respiring and sporulating but not in fermenting diploid cells (Figure 3C) and only poorly transcribed in cell-cycle synchronised haploid cells (data not shown). Moreover, the viewer indicated Ykl162c-a transcription footprint spans only *YKL162C-A* and not the entire ancestral *PIR6* locus. Finally, ribosome profiling (Brar et al., 2012) suggested such transcripts might also be translated (data not shown). Together, the newly-evolved signal sequence and the finely-tuned transcription response suggest that Ykl162c-a might be positively selected for, despite currently having no assigned function.

### 3.4. Hsp150-fusion proteins are well-suited for N-terminally-anchored surface display

After investigating the diversity and 3D structure of *S. cerevisiae* Pir proteins, we used our insights to engineer novel Hsp150 variants, aiming to enhance the N-terminally-anchored surface display. We limited ourselves to rationally redesigning Hsp150 (Pir2), an archetypal Pir protein with 11 repeats. To prevent Pir-specific tandem DNA repeats from recombining, we *de novo* synthesised the gene encoding *S. cerevisiae* Hsp150, thus optimising its codon usage and avoiding repetitive DNA sequences.

To determine whether the *de novo* synthesised Hsp150 fusion proteins can successfully traverse the secretory pathway and anchor themselves to the cell wall, we constructed a strain constitutively expressing a full-length Hsp150 linked to a fluorescent protein, and imaged it with confocal microscopy. For this purpose, we tethered a red fluorescent protein ymScarletI to the Hsp150 C-terminus via a 54 amino acid-long disordered linker encoding three tandem HA-tags and a His-tag (Figure 4A). We placed the construct encoding this fusion protein (Hsp150-ymScarletI) under the strong constitutive *TEF1* promoter and integrated it in the BY 4741 *his3Δ1* locus. Next, we stained the cells with FITC-conjugated concanavalin A, which binds to the mannoprotein layer and thus outlines the cell wall, and imaged the cells by confocal microscopy. In contrast to the exclusively green-fluorescing wild-type cells (Figure 4B), the Hsp150-ymScarletI cells fluoresced in both green and red (Figure 4C), indicating ymScarletI folded successfully. The Hsp150-ymScarletI was detected throughout the cells nonuniformly, mostly in brightly-lit globules, suggesting the fusion protein was contained in the endoplasmic reticulum and transported towards the cell periphery, with a small proportion of Hsp150-ymScarletI localising adjacent to the cell wall, as seen by the faint red ring on the inner border of the mannoprotein layer, indicating the fusion protein was successfully targeted to the cell wall. The faint cell wall-adjacent signal implied that either extracellular conditions dimmed the fluorescence of the fusion protein or that its overexpression had saturated the exporting capacity of the secretory pathway.

**Figure 4:**
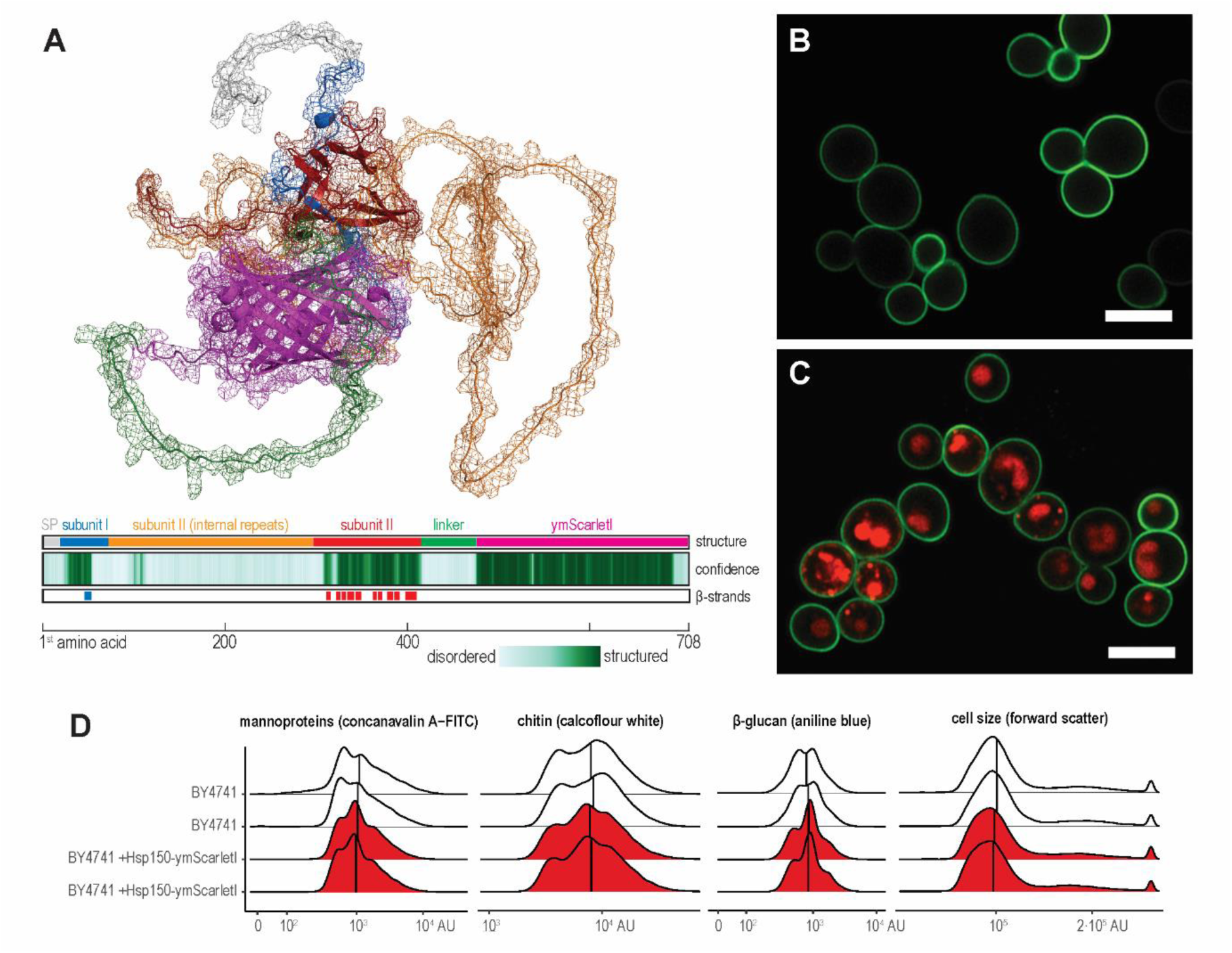
Saturating secretory pathway with Hsp150-ymScarletI fusion protein does not affect the cell wall. A) Alphafold2-structural model of the Hsp150-ymScarletI fusion protein. The colour scheme of the structural model and the summary tracks follows Figure 1. B) Confocal micrograph of the wild-type cells stained with FITC-conjugated concanavalin A. Scale bar denotes 5.0 μm. C) Confocal micrograph of Hsp150-ymScarletI-expressing cells stained with FITC-conjugated concanavalin A. D) Cell size and the amounts of mannoprotein, chitin, and β-glucan of wild-type (white) and Hsp150-ymScarletI-expressing cells (red), quantified via flow cytometry and FITC-conjugated concanavalin A-, calcofluor white-, and aniline blue-staining, respectively. Graphs show two biological replicates.

We wondered whether such saturation of secretory pathway hampered surface display by triggering cell wall remodelling, which could lower the amount of β-glucan available for mannoprotein binding. To test this hypothesis, we quantified three main components of the yeast cell wall via flow cytometry. We stained wild-type and Hsp150-ymScarletI-overexpressing cells with FITC-conjugated concanavalin A, calcofluor white, and aniline blue, which specifically bound to mannoproteins, chitin, and β-glucan, respectively. However, in all three cases, the distributions of the cell wall components between wild-type and Hsp150-ymScarletI cells remained similar (Figure 4D), as did the cell size, suggesting overexpressing Hsp150-ymScarletI did not trigger cell wall stress or remodelling. Thus, strong expression of Hsp150-based fusion proteins does not hamper surface display.

### 3.5. Positioning of the Hsp150’s fusion partner alters the efficiency of the surface display

Next, we measured the efficiency of the Hsp150-based surface display. We constructed an Hsp150-β-lactamase fusion protein and quantified its presence on the cell surface via a colourimetric assay based on nitrocefin, a cephalosporin derivative that does not cross the cell membrane (O’Callaghan et al., 1972; Simonen et al., 1994). In this assay, β-lactamase cleaves nitrocefin’s β-lactam ring, shifting its peak absorbance from 390 nm to 482 nm. Using this assay, we tested five Hsp150-β-lactamase fusion proteins, in which β-lactamase was inserted at the beginning of the Hsp150’s subunit I (v1), after its first internal Pir repeat (v2), at the beginning of the well-structured part of subunit II (v3), immediately after the Hsp150’s C-terminus (v4), or after the disordered tag-rich linker (v5) (Figure 5A).

**Figure 5:**
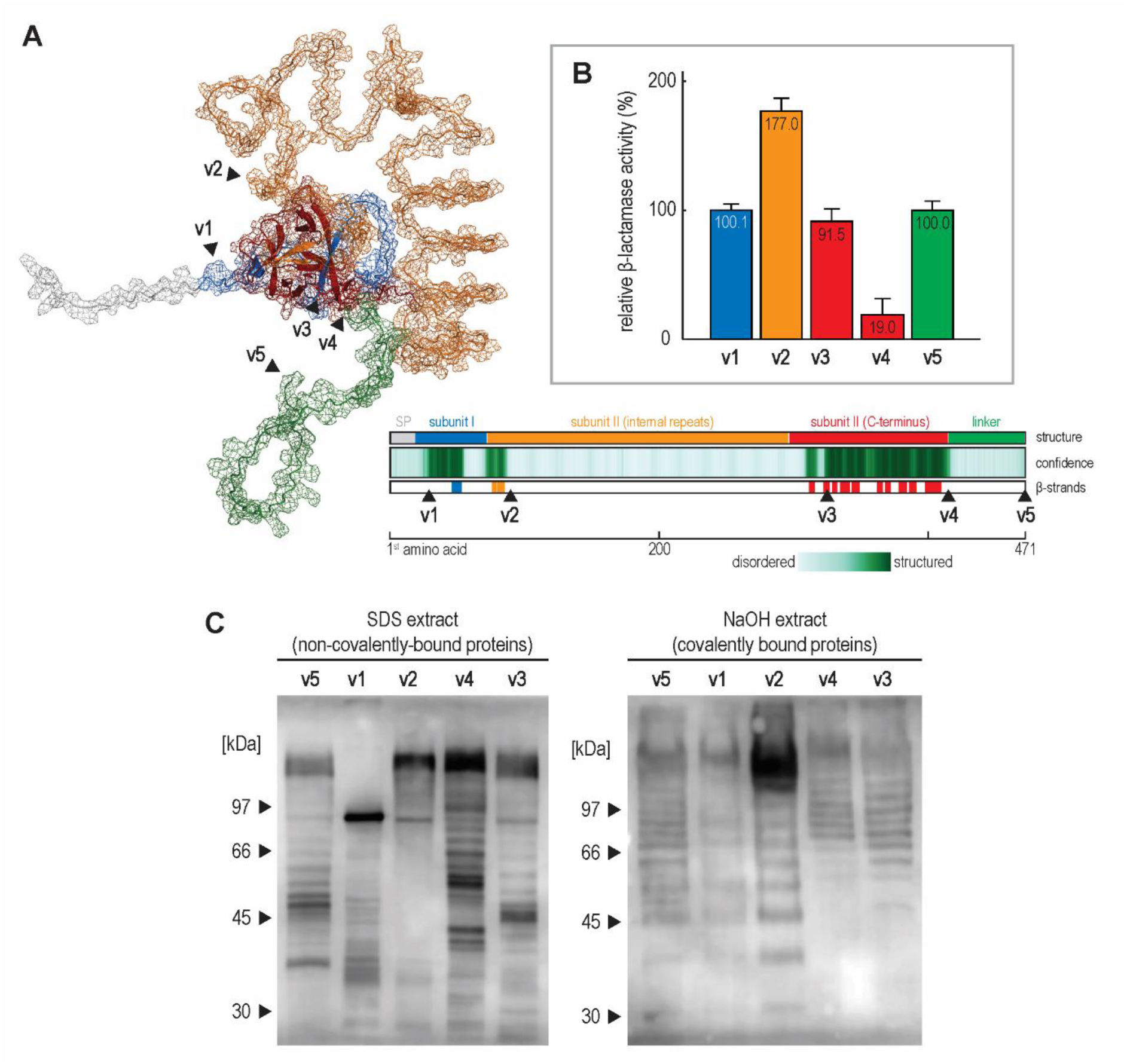
Shuffling the β-lactamase within the Hsp150-fusion protein affects surface display. A) Alphafold2-structural model of Hsp150 protein fused with the tag-rich disordered linker. Black triangles point to the β-lactamase insertion sites. The colour scheme of the structural model and the summary tracks follow Figure 1. B) Relative β-lactamase activity of fusion constructs v1, v2, v3, v4, and v5, normalised to that of construct v5, with error bars denoting standard deviations and bar colours indicating the position of the inserted β-lactamase within the structural model. C) Anti-HA immunoblotting of the noncovalently- and covalently-bound cell wall proteins.

While all five fusion constructs were functional, shuffling the β-lactamase had a striking effect, with constructs differing up to 9-fold in their activity (Figure 5B). Construct v2, carrying β-lactamase after the first internal repeat, performed the best, while construct v4, carrying β-lactamase immediately after Hsp150 C-terminus, performed the worst, reaching only 11% of the activity of construct v2. Interestingly, constructs v1, v3, and v5, carrying β-lactamase at the beginning of subunit I, i.e., immediately after the signal sequence, at the beginning of the well-structured part of subunit II, and after the disordered tag-rich linker, respectively, performed equivalently, reaching 55% of the activity of construct v2. Thus, the choice of β-lactamase insertion site affected the Hsp150-based surface display profoundly.

We then tested whether Hsp150-β-lactamase constructs still bind to the cell surface, covalently via their internal repeats and noncovalently through steric interactions with other cell wall components (Figure 5C). The anti-HA immunoblotting of different fractions of cell wall proteins, targeting the tag-carrying disordered linker, showed that all five constructs bonded to the cell wall covalently and noncovalently, with the most active construct v2 also producing the strongest signal in the covalently-bound fraction. Thus, shuffling β-lactamase within the Hsp150 construct does not preclude covalent and non-covalent binding of the Hsp150-fusion proteins to the cell wall.

The immunoblotting also uncovered an anomalously low molecular mass of noncovalently-bound construct v1, suggesting this construct was cleaved. Indeed, as is the original Hsp150, all five Hsp150-β-lactamase constructs are substrates for Kex2 (Russo et al., 1992), which cleaves them into two polypeptide chains, by hydrolysing the peptide bond between subunit I and II. However, as only the construct v1 carries 29 kDa β-lactamase within subunit I, before the Kex2 processing site, its cleavage is particularly obvious. Interestingly, the non-Kex2-processed form appears only in the covalently-bound fraction, suggesting it more often binds covalently to the cell wall.

The immunoblots also pointed to pronounced yet reproducible and regular degradation of Hsp150-fusion proteins. Such degradation was also observed when detecting chromosomally-tagged Hsp150 and was unresponsive to protease inhibitors (data not shown). With the number of minor bands mostly matching the number of the internal repeats, the signal points either to uncontrolled proteolytic cleavage during protein isolation or a controlled processing event, either of which likely occurs at the unstructured and thus exposed turns of the β-helix.

### 3.6. Identifying minimal Hsp150 region needed for N-terminally anchored surface display

Attempting to design the Pir-tag, i.e., a short peptide sequence that would allow for N-terminally anchored surface display of any protein, we initially constructed six truncated versions of construct v5, lacking various combinations of subunits I and II (Figure 6A). The constructs varied in their display efficiency, which ranged from 2% for construct Δ5, lacking the entire subunit I and the N-terminal part of subunit II, including all internal repeats, to 243% for construct Δ1, lacking only the C-terminal part of subunit II (Figure 6B). The comparison of constructs indicated that the C-terminal part of subunit II strongly impaired surface display, lowering it 1.9-fold (Δ2 vs Δ4), 2.4-fold (v5 vs Δ1) and 29.5-fold (Δ3 vs Δ5). Conversely, the presence of internal repeats promoted surface display. Accordingly, removing the entire subunit II except one Pir repeat from construct v2, which produced construct v6, lowered its surface display efficiency from 177% to 137%. Finally, to test whether the Hsp150 signal sequence acted as a bottleneck, as Hsp150 is secreted non-conventionally (Fatal et al., 2004; Fatal et al., 2002; Grieve and Rabouille, 2011), we exchanged the Hsp150 signal sequence with that of the Ccw12 cell wall protein. However, both the original construct v5 and the derived construct v7 were displayed equally efficiently. Thus, the Hsp150 C-terminal part of subunit II hinders, while its internal repeats promote surface display.

**Figure 6:**
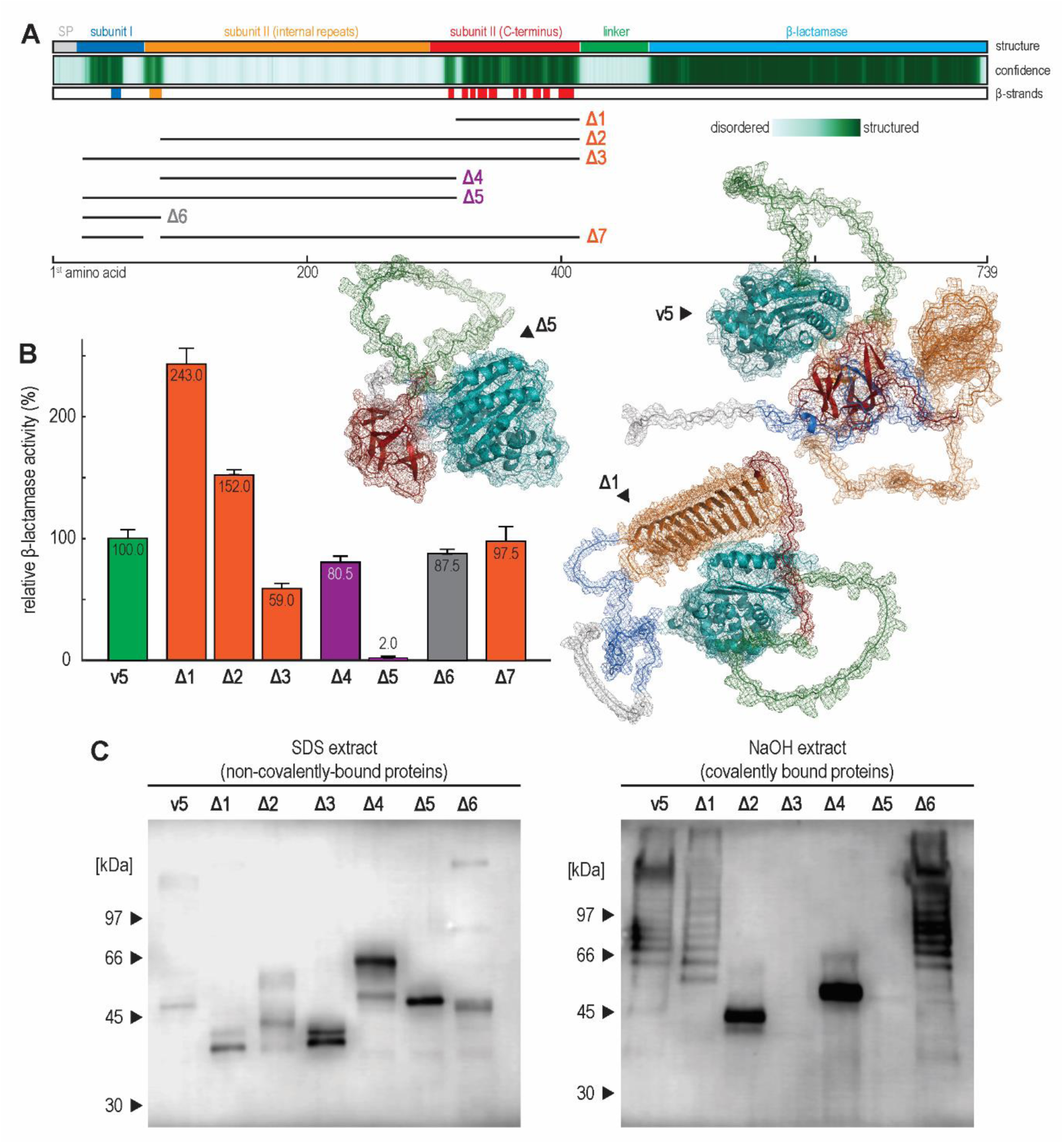
Pinpointing minimal Hsp150 region needed for efficient N-terminally anchored surface display. A) Summary tracks describing construct v5 and deletions introduced into constructs Δ1, Δ2, Δ3, Δ4, Δ5, Δ6 and Δ7, with deleted regions marked as black lines, and the visualisations of Alphafold2-structural models comparing constructs v5, Δ1, and Δ5. B) Relative β-lactamase activity of fusion constructs v5, Δ1, Δ2, Δ3, Δ4, Δ5, Δ6 and Δ7, normalised to that of construct v5, with error bars denoting standard deviations and bar colours corresponding to those in panel A. C) Anti-HA immunoblotting of the noncovalently- and covalently-bound cell wall proteins.

We also tested whether truncated constructs still bind to the cell surface (Figure 6C). The anti-HA immunoblotting showed all constructs bind to the cell wall covalently and noncovalently, save for the Δ3 and Δ5, which, lacking all internal repeats, bind only noncovalently. In line with the previous immunoblotting observations, only constructs with an array of internal repeats produced ladder-like degradation products. Finally, construct Δ3, despite being bound noncovalently, retained significant β-lactamase activity, demonstrating that covalent binding is not indispensable for surface display.

Finally, to produce a minimal Pir-tag, we designed construct Δ7, which lacked the entire Hsp150 subunit I and most of its subunit II, thus preceding the HA-linker and β-lactamase only with a signal peptide and one Pir repeat. Intriguingly, this 42-amino-acid peptide displayed β-lactamase as efficiently as the full, 413-residues long Hsp150, while binding the displayed protein both noncovalently and covalently to the cell wall (data not shown). Thus, construct Δ7 defines a minimal, Hsp150-based peptide for efficient N-terminally anchored surface display.

## 4. Discussion

In this work, we advance yeast surface display by investigating Pir proteins, combining genomic, structural and evolutionary insights to optimally position a protein of interest within *S. cerevisiae* Hsp150. Moreover, we distil the Hsp150 salient features into a minimal Pir-tag, thus allowing for an efficient and easily implementable N-terminally anchored surface display.

Our structural insights, obtained by comparing Alphafold2 predictions with experimentally determined protein structures, suggested that each Pir protein encodes one lectin-like domain. Such an approach could be expanded to other proteins localised to the cell wall, as many still lack clearly-defined functions, being highly and heterogeneously glycosylated and thus challenging to crystallise (Orlean, 2012). Such studies would also hint at the potential physiological roles of cell wall proteins, as most of their single-gene deletion mutants are in standard phenotypic assays indistinguishable from the wild type, owing to their high functional redundancy (Teparić et al., 2020).

As they carry lectin-like domains, Pir proteins could bind yeast cells to the sugars displayed on the surfaces of other microorganisms, plants, or animals. By binding to other microorganisms, *S. cerevisiae* could boost the lethality of its environment-sterilising make-accumulate-consume lifestyle (Hagman et al., 2013), i.e., it could attach itself to its unicellular competitors and selectively augment local ethanol concentration beyond tolerable levels. Conversely, by binding to the plant cell wall, yeast cells could fix themselves to an abundant food source while increasing their wildlife-assisted dispersal. Finally, lectin-like proteins could help yeast to colonise the animal digestive tract, e.g., by aiding overwintering in the wasp gut, which acts as an *S. cerevisiae* reservoir (Stefanini et al., 2012), or by populating the mammalian gut. The latter hypothesis is enticing, as Hsp150 (heat shock protein 150) is induced 7-fold at 37 °C but not required for *S. cerevisiae* thermotolerance (Russo et al., 1992) and was twice as abundant in clinical isolates of *S. cerevisiae*, when compared to the standard laboratory strains (Hsu et al., 2015). Moreover, being modelled as knotted, Pir proteins should fare better in harsher conditions than their unknotted counterparts and thus perform well in the gut of warm-blooded animals (Sułkowska et al., 2008; Taylor, 2007). As standard laboratory tests do not assess interspecies dynamics, such an ecological role would also explain the perceived lack of phenotype in single Pir deletion mutants.

With the cell wall being the primary interface between the microbial cell and the environment (Teparić et al., 2020), the sugar-binding character of Pir proteins likely drives their maintenance, diversification, and subfunctionalisation. Indeed, Lozančić et al. (2021) detected Pir proteins in each of the 78 ascomycete yeasts most closely related to *S. cerevisiae*, with several of these species encoding over a dozen of Pir proteins. Moreover, while lectin-like *S. cerevisiae* Pir domains consist of a well-conserved β-stranded core, their interstrand turns, which determine sugar-binding specificity, differ. The tendency toward subfunctionalisation also explains why *S. cerevisiae* would retain both Pir loci after the whole genome duplication and why Pir proteins are differentially expressed throughout the cell cycle and growth phases (Lardenois et al., 2011). Finally, it is interesting to speculate whether the loss of Pir6 and the evolution of Ykl162c-a was positively selected for by the domestication of *S. cerevisiae*, which affected its growth and life cycle remarkably (De Chiara et al., 2022).

The Ykl162c-a is an interesting peptide, encoding only 50 amino acid residues, 21 of which form a newly evolved functional signal sequence. The peptide also avoids all four C-terminal cysteine residues conserved across the Pir family (Toh-E et al., 1993). Moreover, Ykl162c-a evolved a novel transcriptional footprint, as only its open reading frame is transcribed, while the rest of the ancestral Pir6 locus remains silent. While the here-presented results do not confirm that the native transcript is translated and secreted, the putative secreted peptide could act as an (interspecies) pheromone or even an antimicrobial molecule (Rebuffat, 2022). Its presence also suggests that other such peptides with high secretion potential could be encoded yet overlooked within the yeast genome.

The hallmarks of the Pir proteins are their eponymous tandem repeats, whose purpose currently remains unknown. The here-presented models of Pirs as knotted proteins partly discredit the simplest hypothesis, that the repeats simply tether the C-terminal part of the protein far from the cell’s surface. An alternative hypothesis is that Pir proteins serve as stretchable proteinaceous cell wall netting covalently linking β-glucans and thus helping to stabilise the cell wall during growth and remodelling (Orlean, 2012). That hypothesis seeks to explain why the deletion of multiple Pir proteins weakens the cell wall by presuming many Pir proteins bind to β-1,3-glucan through two or more Pir repeats. However, our modelling suggests a third possibility, that by folding into a two-faced β-helix, Pir repeats build a structure with which the cell actively modulates its environment, as do other such structures found in biofilm matrices, adhesins, and antifreeze proteins (DeBenedictis and Keten, 2019).

Our efforts to advance N-terminally anchored surface display focused on *S. cerevisiae* Hsp150, an archetypal and well-investigated member of the Pir protein family (Makarow et al., 2006; Simonen et al., 1994). However, although this protein was previously employed for surface display, a meticulous and called-for benchmarking of its potential insertion sites remained unrealised (Yang et al., 2014), partly because a proper comparison would need to rely on the previously unknown Hsp150 structure.

To rigorously quantify the Hsp150-based surface display, we implemented several strategies. We fused Hsp150 with a β-lactamase reporter protein and employed a nitrocefin assay, which relies on nitrocefin’s cell-membrane impenetrability. Moreover, we placed the Hsp150-fusion protein under the strong inducible *PHO5* promoter (Korber and Barbaric, 2014), ensuring that during preculture, the Hsp150-constructs did not interfere with cell growth. Finally, with confocal microscopy and flow cytometry, we verified at the single-cell and population levels that the strongly expressed fusion protein did not perturb cell wall homeostasis, i.e., did not change the amounts of mannoproteins, chitin, and β-glucans in the cell wall.

By inserting β-lactamase at varying positions throughout Hsp150, we demonstrated that the C-terminal Pir domain inhibits surface display. Thus, inserting β-lactamase next to the Hsp150 C-terminus, without separating the reporter from the terminus with the unstructured HA-spacer, eroded β-lactamase activity. However, the β-lactamase remained unimpaired when inserted just before the C-terminal domain, the position at which the reporter would fold before the C-terminal Hsp150 domain would. As immunoblotted constructs produced equally strong signals, despite having a 5-fold difference in activity, impaired surface display probably stemmed from improper protein folding and not degradation of the fusion protein in the secretory pathway. Conversely, inserting β-lactamase in the long unstructured region, immediately after the first Pir repeat, helped it to fold and possibly helped Hsp150 to bind to the cell wall covalently, as evidenced by the strong immunoblot signal.

The immunoblot of construct v1 also suggested that non-Kex2-processed Hsp150-fusion proteins covalently bind to the cell wall more often than Kex2-processed forms, which was not noted previously. This result also demonstrated that covalent binding to the cell wall is not necessary for surface display, as smaller Kex2-cleaved subunit retained nominal β-lactamase activity, despite lacking Pir repeats. Our future experiments will focus on this and other potential Pir processing events, which were hinted at by the appearance of a ladder-like immunoblot signal.

Construct truncations also indicated that the C-terminal Hsp150 domain undermined while the unstructured repeat-rich region promoted the efficiency of the surface display. Thus, the constructs without the C-terminal domain consistently performed better, especially if the domain comprised most of the construct, as did the constructs containing the repeat region, possibly because cell wall proteins could require unstructured regions to pass through the inner polysaccharide layer efficiently, as was recently shown for bacterial cell wall proteins (Halladin et al., 2021). The C-terminal domain and the repeat-region effects were additive, as the best-performing construct lacked the C-terminal domain but carried the entire subunit I and all repeats. Thus, the Hsp150-derived portion of such a construct promoted a 2.5-fold more efficient surface display than the full-length Hsp150. Moreover, through extensive cropping, we developed a Pir-tag, as efficient as full-length Hsp150 but spanning only 4.5 kDa, i.e., 45 amino acids, 18 of which encode a signal sequence. These two constructs are a much-needed addition to the yeast surface display toolbox that will allow for the efficient and routine refitting of any protein into its N-terminally anchored surface-displayed isoform.

## 5. Conclusion

In this work, we leveraged the structural and evolutionary insights into Pir proteins to augment the N-terminally anchored yeast surface display. With this strategy, we designed two constructs, one 2.5-fold more efficient than the wild-type Hsp150 and another as efficient as the full-length Hsp150 but spanning only 4.5 kDa. As such, we outline a blueprint for efficiently refitting any protein for N-terminally-anchored display on the outer yeast surface.

## Acknowledgements

We thank David Gosset for his help with fluorescent microscopy and flow cytometry on the MO2VING platform (Centre de Biophysique Moléculaire, CNRS, Orleans). Graphical abstract was created with BioRender.com.

## Supplementary material

Details of the plasmid and strain construction.

## Competing Interests

The authors declare no competing interests associated with the manuscript.

## Funding

This research was funded by The Croatian Science Foundation, grants No. IP-2019-04-2891 and DOK-2021-02-9672.

## CRediT author statement

**Tea Martinić Cezar:** Methodology, Investigation, Formal analysis. **Mateja Lozančić:** Methodology, Investigation, Formal analysis. **Ana Novačić:** Methodology, Investigation, Formal analysis, Writing - Review & Editing, **Ana Matičević:** Investigation. **Dominik Matijević:** Investigation. **Béatrice Vallée:** Resources, Supervision. **Vladimir Mrša:** Resources, Supervision, Project administration, Funding acquisition. **Renata Teparić:** Methodology, Resources, Supervision, Writing - Review & Editing, Project administration, Funding acquisition. **Bojan Žunar:** Conceptualization, Methodology, Software, Validation, Investigation, Formal analysis, Visualization, Writing - Original Draft, Writing - Review & Editing.

## Notes

### Competing Interest Statement

The authors have declared no competing interest.

